# Psychiatric risk gene *NT5C2* regulates protein translation in human neural progenitor cells

**DOI:** 10.1101/468546

**Authors:** Rodrigo R.R. Duarte, Nathaniel D. Bachtel, Marie-Caroline Côtel, Sang H. Lee, Sashika Selvackadunco, Iain A. Watson, Gary A. Hovsepian, Claire Troakes, Gerome D. Breen, Douglas F. Nixon, Robin M. Murray, Nicholas J. Bray, Ioannis Eleftherianos, Anthony C. Vernon, Timothy R. Powell, Deepak P. Srivastava

**Affiliations:** MRC Social, Genetic & Developmental Psychiatry Centre, Institute of Psychiatry, Psychology & Neuroscience, King’s College London, London, United Kingdom.; Department of Basic & Clinical Neuroscience, Institute of Psychiatry, Psychology & Neuroscience, King’s College London, London, United Kingdom.; Department of Biological Sciences, Columbian College of Arts and Sciences, The George Washington University, Washington, District of Columbia, United States.; MRC Centre for Neurodevelopmental Disorders, King’s College London, London, United Kingdom.; MRC London Neurodegenerative Diseases Brain Bank, Department of Basic & Clinical Neuroscience, Institute of Psychiatry, Psychology & Neuroscience, King’s College London, London, United Kingdom.; Division of Infectious Diseases, Weill Cornell Medicine, Cornell University, New York, New York, United States.; Department of Psychosis Studies, Institute of Psychiatry, Psychology & Neuroscience, King’s College London, London, United Kingdom.; MRC Centre for Neuropsychiatric Genetics and Genomics, Cardiff University School of Medicine, Cardiff, United Kingdom.

**Author notes:** Correspondence to: Dr Deepak P Srivastava, Maurice Wohl Clinical Neuroscience Institute, Institute of Psychiatry, Psychology & Neuroscience, King’s College London, 5 Cutcombe Road, London SE5 9RX, United Kingdom. Shared senior authors (Timothy R. Powell, Deepak P. Srivastava).

**Keywords:** 5’ AMP-activated protein kinase (AMPK), rpS6, drosophila melanogaster, postmortem tissue, psychiatric disorders, GWAS, functional genetics, fruit fly, human neural progenitor cells, neurodevelopment

## Abstract

Genome-wide significant variants associated with combined risk for major psychiatric disorders on chromosome 10q24 affect the expression of the cytosolic 5’-nucleotidase II (*NT5C2, cN-II*) in population controls, implicating it as a psychiatric susceptibility gene. Risk alleles are associated with reduced expression of this gene in the developing and adult brain, but the resulting neurobiological risk mechanisms remain elusive. In this study, we provide further evidence for the association of *NT5C2* with psychiatric disorders, and use a functional genetics approach to gain a deeper understanding of the function of this risk gene in the nervous system. *NT5C2* expression was significantly reduced in the *post-mortem* brain of schizophrenia and bipolar disorder patients, and its protein predominately expressed in neurons within the adult brain. Using human neural progenitor cells (hNPCs), we found that *NT5C2* expression peaked at the neural progenitor state, where the encoded protein was ubiquitously distributed through the cell. *NT5C2* knockdown in hNPCs elicited transcriptomic changes associated with protein translation, that were accompanied by regulation of adenosine monophosphate-activated protein kinase (AMPK) signalling and ribosomal protein S6 (rpS6) activity. To identify the effect of reduced neuronal *NT5C2* expression at a systems level, we knockdown its homologue, *CG32549*, in *Drosophila melanogaster* CNS. This elicited impaired climbing behaviour in the model organism. Collectively, our data implicate *NT5C2* expression in risk for psychiatric disorders and in *Drosophila melanogaster* motility, and further suggest that risk is mediated via regulation of AMPK signalling and protein translation during early neurodevelopment.

## Introduction

Genetic variants on chromosome 10q24 are associated with combined risk for schizophrenia, bipolar disorder, major depression, autism, and attention deficit hyperactivity disorder^1^, and constitute the third top association signal in the latest schizophrenia genome-wide association study (GWAS)^2^. Previous work from our group has demonstrated that the psychiatric risk variants at this locus exert *cis-*regulatory effects on the cytosolic 5’-nucleotidase II gene (*NT5C2, cN-II*) in population controls, reducing expression of this gene in the adult and developing brain^3^. However, the neurobiological mechanisms through which genetic variation at the *NT5C2* locus may increase risk for psychiatric conditions remain elusive. Interestingly, *NT5C2* has also been implicated in a myriad of other medical conditions, including intellectual disability^4^, Parkinson’s disease^5^, spastic paraplegia^6^, cardiovascular disease^7, 8^, and acute lymphoblastic leukaemia^9^, which suggests that *NT5C2* may regulate fundamental aspects of cell biology that are ultimately dysregulated in these conditions.

The *NT5C2* gene produces an enzyme that cleaves inorganic phosphate from purine and purine-derived nucleotides such as adenosine, inosine and guanosine monophosphate (AMP, IMP, and GMP, respectively)^10^, or that catalyses the transfer of phosphate groups between purine nucleotides and nucleosides^11^. Purinergic compounds have been shown to regulate cell cycle progression and to act as neurotransmitters, and neurotrophic or neuroprotective agents^12, 13^, which is unsurprising given their role in fundamental metabolic processes such as DNA replication, gene transcription and protein synthesis^14, 15^. Interestingly, a purinergic hypothesis of schizophrenia has been proposed, which explains neurodevelopmental and neurochemical aspects of this disorder^16^. One plausible mechanism via which NT5C2 may regulate such pivotal processes is via *adenosine monophosphate-activated protein kinase* (AMPK) signalling, which is a major regulator of cellular energy homeostasis^17-19^. It remains unclear, however, whether this mechanism occurs in the context of the nervous system, or the cell types involved.

The present work provides the most in-depth characterisation of the distribution, expression and function of *NT5C2* in the brain and in cultures of human neural progenitor cells (hNPCs). First, to confirm that *NT5C2* is a psychiatric risk gene, we explore its expression in RNA sequencing (RNA-seq) data from the brain of schizophrenia, major depression and bipolar disorder patients. This analysis revealed that *NT5C2* expression is significantly reduced in schizophrenia and bipolar disorder patients relative to unaffected controls. We also used *post-mortem* human brain tissue to identify the major cell types expressing NT5C2 in the adult brain, revealing this protein is more expressed in neurons relative to glial cells. To gain insight into the molecular mechanisms that this gene influences, we investigated the expression, sub-cellular distribution and function of *NT5C2* in hNPCs. This revealed that the gene is highly expressed during neurodevelopment and that the NT5C2 protein is ubiquitously distributed in the soma and cellular processes of neural progenitors. Using an RNA interference-mediated approach (RNAi), reduced expression of *NT5C2* in hNPCs elicited transcriptomic changes associated with protein translation regulation, which were accompanied by differential regulation of AMPK signalling and ribosomal protein S6 (rpS6) activity. Finally, to elucidate the impact of reduced *NT5C2* expression at a systems level, we utilised a *Drosophila melanogaster* (*D. melanogaster*) model in which the fly homologue of *NT5C2*, *CG32549*, was knocked-down either ubiquitously or specifically within the nervous system. Collectively, our work describes the hitherto unknown pattern of *NT5C2* distribution and expression in the adult brain and hNPCs, implicating it in the regulation of AMPK signalling and protein translation.

## Materials and Methods

See Supplementary Information for further details on Materials and Methods.

### Brain samples

To identify cell-type specific expression of *NT5C2* in the adult brain, we obtained samples from unaffected controls from the Medical Research Council London Neurodegenerative Disease Brain Bank, at the Institute of Psychiatry, Psychology & Neuroscience, King’s College London, under the license of the United Kingdom Human Tissue Authority (ref. 12293). Demographics of the *post-mortem* samples is available in the Supplementary Information.

### RNA-sequencing analysis

To study expression of *NT5C2* in the brain of psychiatric patients, we analysed RNA-seq data from the Stanley Neuropathology Consortium^20^. This cohort consisted of a collection of matched hippocampus samples from subjects diagnosed with bipolar disorder, schizophrenia, or major depression, and unaffected controls (n = 15 each). Reads were trimmed using Trimmomatic 0.36^21^, and mapped to Ensembl genes (build 38, v93) using kallisto^22^. Differential expression was calculated using Wald tests in DESeq2^23^ whilst controlling for the effect of demographics, and corrected using a false discovery rate (FDR) cut-off of 10% (q < .10). Raw and normalised counts, and factors and covariates considered in the analysis are available in **Supplementary Tables 1-3**. Raw RNA-seq data is available upon request via http://sncid.stanleyresearch.org.

### Immunohisto- and cytochemistry

We used immunolabelling to quantify the *NT5C2* knockdown in hNPCs and identify its sub-cellular distribution in hNPCs and brain tissue. Brain sections were deparaffinised and submitted to antigen retrieval and autofluorescence removal protocols (please see Supplementary Information for more information). Brain sections and cell cultures were permeabilised and blocked and incubated with the following primary antibodies: NT5C2 (M02-3C1) (Abnova, Taipei, Taiwan), IBA1 (Menarini Diagnostics, Winnersh, United Kingdom), GFAP (Dako Agilent, Santa Clara, United States), MAP2, Parvalbumin and Beta-III-Tubulin (Abcam, Cambridge, United Kingdom). Fluorescently labelled secondary antibodies included Goat Alexa 488, 568 or 633 antibodies (Thermo Fisher Scientific).

### Cell lines

To identify the distribution and function of *NT5C2 in vitro*, we analysed hNPCs from the CTX0E16 neural stem cell line^24^ or from human induced pluripotent stem cells (hiPSCs) from an unaffected control^25^, and human embryonic kidney cells 293T (HEK293T). The CTX0E16 neural cell line^24^ was obtained from ReNeuron Ltd. under a Material Transfer Agreement. Cells were derived and maintained as described in the Supplementary Information and elsewhere^24, 25^.

### RNA and protein isolation and quantification

To identify gene and protein expression and phosphorylation differences associated with *NT5C2* function, we isolated total RNA or protein from *in vitro* cultures using TRI Reagent (Thermo Fisher Scientific, Waltham, Massachusetts, United States) or RIPA Buffer supplemented with Halt Protease and Phosphatase Inhibitor Cocktail (Thermo Fisher Scientific), respectively. Details on reverse transcriptase quantitative polymerase chain reaction (RT-qPCR), quality control and western blotting are available in the Supplementary Information. Primary antibodies for western blotting included: AMPK-alpha (D6) and phospho-AMPK-alpha (Thr172) (Santa Cruz Biotechnology, Dallas, Texas, United States), and total rpS6 (54d2) and phospho-rpS6 (Ser235/Ser236) (Cell Signalling, Danvers, Massachusetts, United States).

### Confocal microscopy

We imaged fluorescently labelled cultures or brain sections using confocal microscopy to identify the distribution of NT5C2 in the adult brain, and the sub-cellular distribution of this protein in hNPCs. Imaging was performed at the Wohl Cellular Imaging Centre, King’s College London, using a Nikon A1R (Nikon, Amsterdam, Netherlands) or a Leica SP5 Confocal Microscope (Leica, Wetzlar, Germany). Images were taken as z-stacks of 8-10 plans, and exported to Fiji, where background subtracted images and maximum intensity projections were generated. To identify cell-type expression of *NT5C2* in the brain, co-localisation was defined as percentage of co-localised clusters relative to total number of clusters detected per image. High throughput analysis was performed using an ImageJ macro^26^ previously used for co-localisation investigations^27^ (n = 4 control subjects, 3 technical replicates per antibody combination, 20 fields of view (FOV) each). A detailed summary of this macro can be found in Supplemental Information. To quantify the knockdown in hNPCs, regions of interest (ROI) were defined based on beta-3-tubulin expression, and NT5C2 corrected total cell fluorescence (CTCF) values were calculated as: *CTCF = integrated density - (area × mean fluorescence of three background readings)* (n = 4 biological replicates per condition, 4 FOV each).

### Microarray analysis

A microarray analysis was performed to characterise the transcriptomic changes associated with the *NT5C2* knockdown in hNPCs. Samples were analysed at the Institute of Psychiatry, Psychology & Neuroscience BRC Genomics & Biomarker Core Facility, King’s College London, using Human HT12 v4 BeadChip arrays (Illumina, Cambridge, Cambridgeshire, UK). A linear regression model was used to quantify gene expression differences between conditions, whilst controlling for the effect of confounders such as biological replicate and microarray batch. The expression data were deposited in GEO under accession code GSE109240. Enrichment for Gene ontology (GO) terms was calculated against the background of all genes in the genome using GeneMania^28^.

### Fly stocks and motility test

To test the role of *NT5C2* in psychomotor behaviour in *D. melanogaster*, the *D. melanogaster* homologue of *NT5C2*, homologue *CG32549*, was knocked down by crossing the CG32549-RNAi line (v30079) with fly lines containing Gal4-driven promoters of genes that are ubiquitously expressed (*ACT5C:* BL4414), neuronal-specific (*ELAV:* BL8765), or gut-specific (*GUT:* DGRC113094). Negative geotaxis was used to calculate climbing success and assess psychomotor behaviour, as previously described^29^. Survivorship was determined 17-20 days post eclosure, as the number of flies alive out of the initial 20 flies allocated per tube.

### Statistical analysis

To identify differences between more than two independent groups, we used one-way ANOVAs followed by Tukey post hoc tests if values were normally distributed (e.g. co-localisation between NT5C2 and different markers, or the effect of two independent siRNAs in hNPCs relative to control cultures); or Kruskal-Wallis tests followed by Dunn’s post hoc tests, if values were not normally distributed (e.g.: the effect of the knockdowns on total and phosphorylated levels of AMPK and rpS6). To compare differences between two groups, we performed t-tests if values were normally distributed (e.g. expression differences between hNPCs and cultures, the effect of the knockdown or overexpression in hNPCs or HEK293T); or Mann-Whitney tests if they were not normally distributed (e.g. survival and climbing success ratios associated with *CG32549* knockdowns in Drosophila); correction for multiple testing was performed using the Bonferroni method. The Fisher’s exact test to calculate significance of the gene overlaps was performed in R using the package ‘GeneOverlap’. Statistical analyses were performed in R or in IBM SPSS.

## Results

### *NT5C2* expression is reduced in the brain of schizophrenia and bipolar disorder patients

We previously demonstrated that *NT5C2* expression is reduced in the brain of unaffected controls due to cis-regulatory effects associated with psychiatric risk alleles located on chromosome 10q24^3^. To study the expression of this gene in a psychiatric cohort, we analysed RNA-sequencing (RNA-seq) data from the hippocampus of patients diagnosed with bipolar disorder (BD), major depression (MDD), or schizophrenia (SCZ), and unaffected controls. We assessed gene expression differences associated with case-control status (**Supplementary Tables 4-6**) whilst controlling for potential confounding effects of demographics (**Supplementary Tables 7-12**). This analysis revealed that *NT5C2* was less expressed in SCZ (P < .001, false discovery rate (FDR) corrected P = .01, fold-change = 0.56) and BD patients (P < .001, corrected P = .02, fold-change = 0.69), but not in MDD patients (P > .05) (**Figure 1A**). These findings corroborate a role for *NT5C2* in mediating susceptibility to psychiatric disorders, particularly those associated with psychotic features.

**Figure 1.**
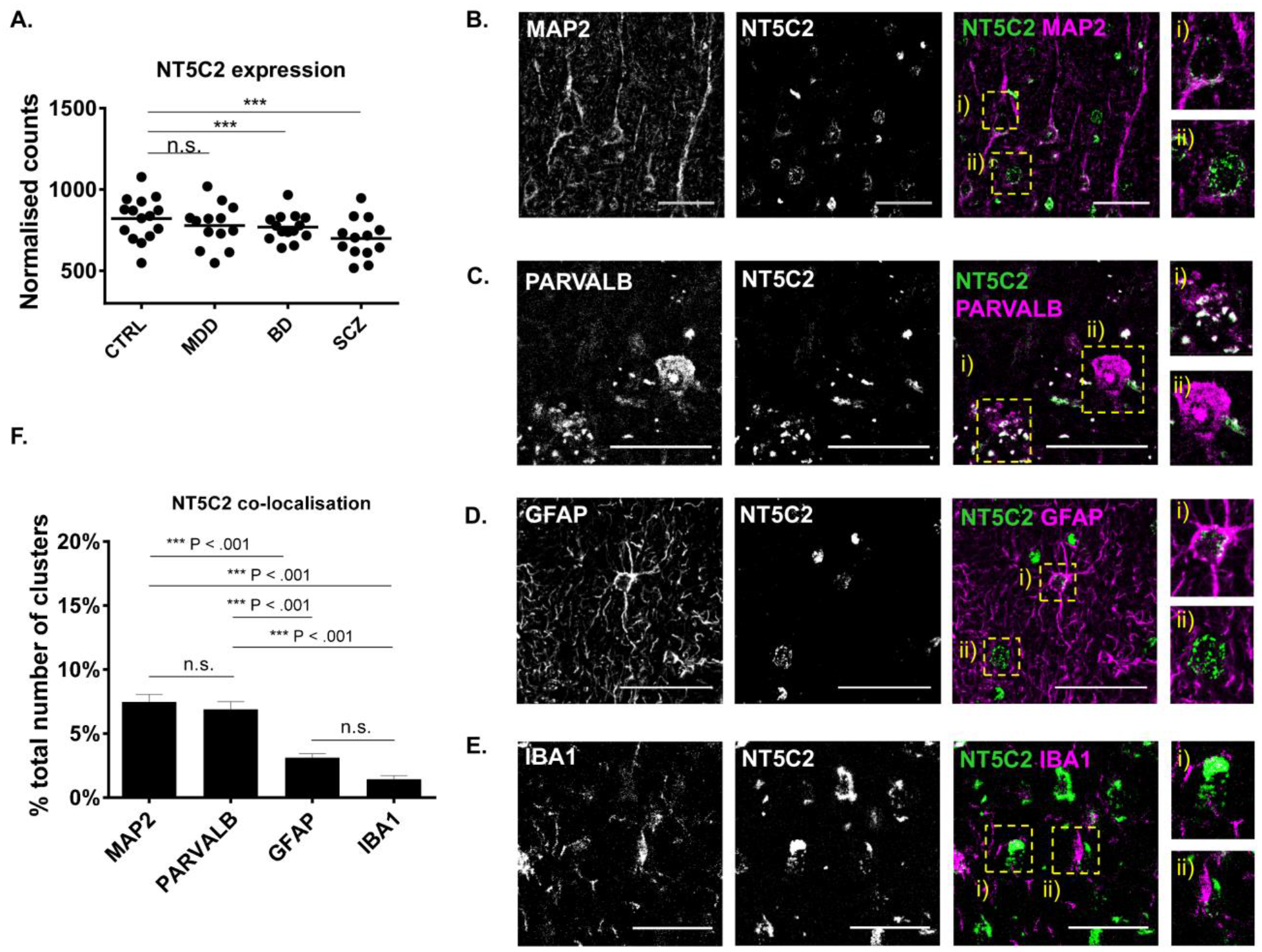
Expression of the psychiatric risk gene in patients, and distribution of the encoded protein in unaffected controls. **(A)** *NT5C2* is downregulated in the hippocampus of bipolar disorder and schizophrenia patients (Wald tests, q < .001) **(B)** Co-localisation of NT5C2 staining with MAP2 (neuronal marker), **(C)** PARVALB (interneuron marker), **(D)** GFAP (glial marker), **(E)** and IBA1 (microglia marker). Scale bars are 50 µM. **(F)** Quantification of the co-localisation of NT5C2 with markers from **B** to **E** revealed a significant difference in the co-localisation of NT5C2 with these markers (one-way ANOVA, P < .001, Tukey post hoc tests: ***P < .001 for all comparisons).

### Expression of NT5C2 is enriched in neurons relative to glial cells in the adult brain

To understand how genetic variation affecting NT5C2 expression may confer risk for psychiatric disorders, we investigated which neuronal cell types expressed this gene. First, we examined single-cell RNA-seq data from the mouse cortex^30^ to predict which cell type(s) highly expressed *NT5C2*. This analysis revealed a significant difference in the cell-type specific expression of *NT5C2* within neuronal and non-neuronal cells (One-way ANOVA, F (3, 1008) = 11.11, P < .001). Post-hoc analysis confirmed that *NT5C2* is more abundant in neurons and interneurons than astrocytes (Tukey post hoc tests: P < .001 for all comparisons; **Supplementary Figure 1**).

To investigate whether this distribution pattern also occurs in humans, we performed a series of immunocolocalisation experiments using autopsy brain tissue and confocal microscopy. We confirmed specificity of an antibody raised against NT5C2 by using it to probe a gain-of-function in human embryonic kidney cells 293T (HEK293T) and CTX0E16 hNPCs (**Supplementary Figures 2 and 4**), and loss-of-function in hNPCs (**Figures 3C and D**). We next analysed the distribution of NT5C2 in the prefrontal cortex of *post-mortem* brain using standard immunoperoxidase staining with DAB as the chromogen (**Supplementary Figure 3**). A qualitative analysis of NT5C2-positive immunostaining with Nissl counter-stain to reveal cellular morphology suggested that NT5C2 was present in neurons, glia and the surrounding neuropil **(Supplementary Figure 3)**. However, we noted that not all putative glial cells expressed NT5C2 (red arrows; **Supplementary Figure 3**). Therefore, to confirm this, we quantified the cell type-specific expression of NT5C2 by measuring the co-localisation of this protein with markers of mature neurons (microtubule-associated protein 2, MAP2), a sub-class of gamma-amino butyric acid (GABA) interneurons (parvalbumin, PARVALB), astrocytes (glial fibrillary acidic protein, GFAP), and microglia (ionized calcium-binding adapter molecule 1, IBA1; **Figures 1B-F**). We included PARVALB on the basis of the wealth of complementary lines of evidence implicating these cells in the pathophysiology of psychiatric disorders^31^. This analysis revealed a significant difference in the specific co-localisation of NT5C2 with each of these markers (One-way ANOVA, F (3,44) = 39.12, P <.001, n = 4 control subjects). Post-hoc analysis confirmed that the mean percentage co-localisation values significantly differed between neuronal and non-neuronal markers (MAP2: 7.48% ± 2.02 (standard deviation); PARVALB: 6.89% ± 2.09; GFAP: 3.13% ± 1.09; IBA1: 1.44% ± 0.93; Tukey post hoc tests: P < .001 for all comparisons; **Figure 1F**), but not within these categories (i.e., GFAP vs. IBA1, P > .05). These data corroborate the single-cell RNA-seq data from the mouse brain (**Supplementary Figure 1**), and confirm our qualitative observations using immunoperoxidase staining (**Supplementary Figure 3**). Overall, these data suggest that whilst NT5C2 expression at the message and protein level is found in both neurons and glial cells, it is clearly enriched in neurons relative to glia. This is consistent with the recent observation that there is an enrichment for expression of psychiatric risk genes in neuronal cells^32^.

### NT5C2 is highly expressed and ubiquitously distributed in hNPCs

The role of *NT5C2* in psychiatric disorders commence during neurodevelopment^3^, a period underscored by a large number of concurrent and ongoing complex process, implicated in all major psychiatric disorders. Indeed, assessment of the Human Brain Transcriptome Atlas^33^ (**Figure 2A**) has identified that *NT5C2* expression peaks during this period. Thus, to investigate *NT5C2* function in neurodevelopment, we explored the sub-cellular distribution and molecular function of this gene in hNPCs. First, we characterise expression of *NT5C2* RefSeq transcripts NM_012229 and NM_001134373 in hNPCs from the CTX0E16 neural progenitor cell line and neurons terminally differentiated for 28 days (DD28)^24^. At this stage, terminally differentiated CTX0E16 cultures mainly comprise of neurons (∼80%) and glial cells (∼10%)^24^. This analysis revealed that both *NT5C2* transcripts were expressed ∼30% higher in hNPCs compared to DD28 neurons on average. Although peak expression was observed in hNPCs, both transcripts displayed persistent expression at moderate to high levels in DD28 cultures (**Figure 2B**), indicating that *NT5C2* may also play a role in immature neurons. These data are consistent with a psychiatric risk mechanism mediated by *NT5C2* expression which starts during neurodevelopment and persists during adult life^3^.

**Figure 2.**
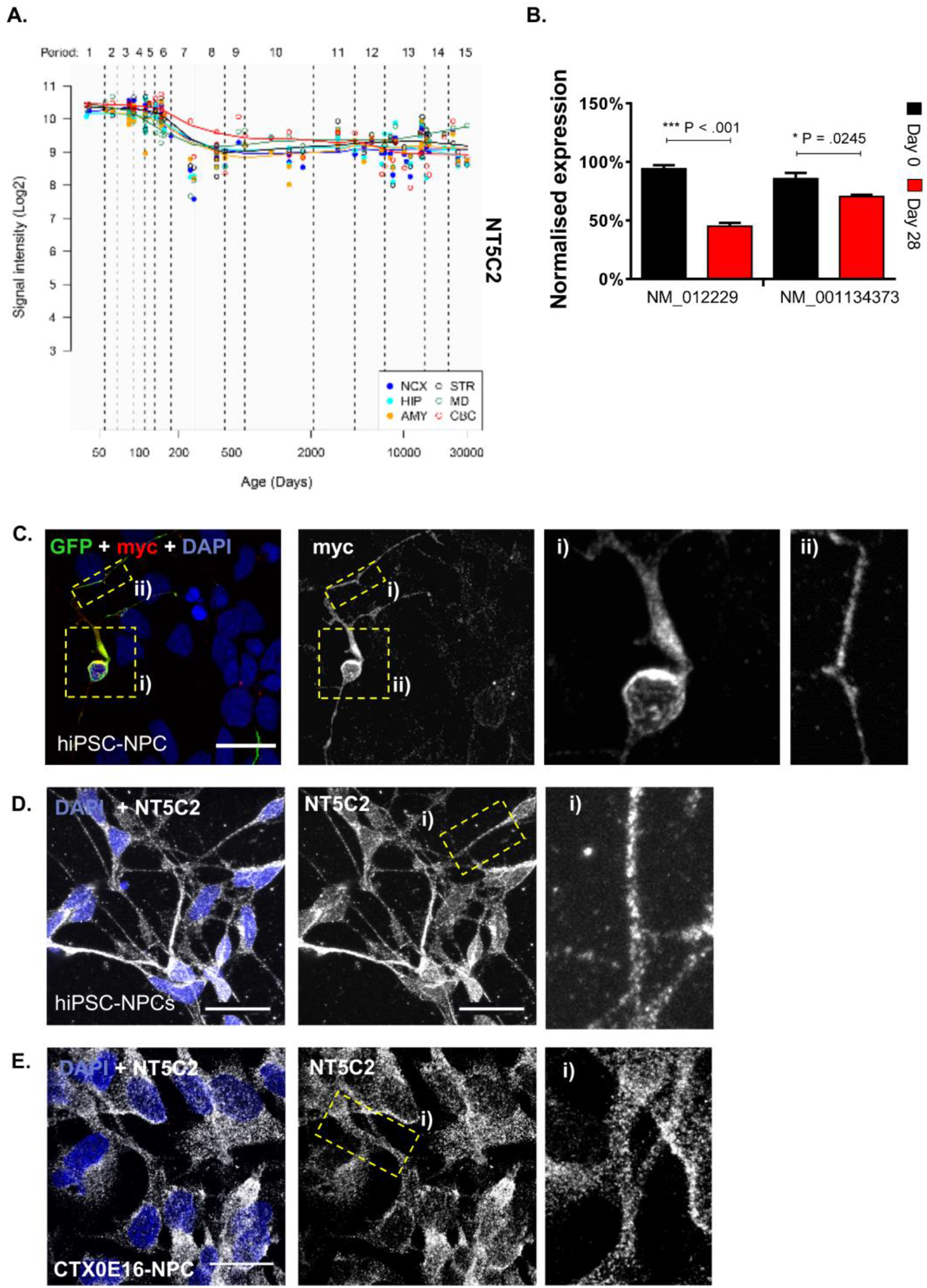
Neurodevelopmental expression of *NT5C2*, and protein distribution in human neural progenitor cells. **(A)** Neural expression of *NT5C2* across human development, according to the Human Brain Transcriptome Atlas^33^, showing that expression peaks during foetal development. **(B)** The expression of *NT5C2* RefSeq transcripts NM_012229 and NM_001134373 in hNPCs (Day 0) and immature neurons differentiated from this source for 28 days. Expression is significantly higher at the neural progenitor state relative to the later stage of differentiation (independent sample t-tests: ***P < .001, *P <.05). **(C)** Distribution of ectopic NT5C2 was assessed by transfecting hNPCs with pNT5C2-myc and peGFP plasmids, followed by immunolabelling using antibodies raised against myc or GFP. GFP expression was used as morphological marker to outline cell morphology. Exogenous protein expression was observed along the cell soma and neurites. **(D)** Subcellular localisation of endogenous NT5C2 in hNPCs derived from a hiPSC line, or **(E)** or from the CTX0E16 cell line. In both hNPC systems, NT5C2 demonstrated a similar distribution pattern, suggesting this protein is ubiquitously distributed in neural progenitor cells. Scale bars are 20 µM.

Next, we assessed the sub-cellular distribution of *NT5C2* in hNPCs derived from human induced pluripotent stem cell (hiPSC) and CTX0E16 cells. First, we ectopically expressed a myc-tagged NT5C2 construct in hiPSC-NPCs. This revealed that ectopic myc-NT5C2 was abundantly expressed in the cell soma and was present in punctate structures along neurites (**Figure 2C**). We further confirmed the ability of our antibody raised against NT5C2 to detect myc-NT5C2 (**Supplementary Figure 4**). Next, we examined the distribution of *endogenous* NT5C2 in hNPCs derived from hiPSCs (**Figure 2D**) and from the CTX0E16 cell line (**Figure 2E**). Similar to the distribution of ectopic protein, endogenous NT5C2 was found dispersed throughout the cell, and was present in punctate structures within the cell soma and along neurites; virtually all imaged cells expressed NT5C2. Taken together, these data suggest that NT5C2 is highly expressed and ubiquitously distributed in hNPCs.

### Reduced NT5C2 expression in hNPCs is associated with regulation of protein translation

In order to further understand how NT5C2 function may impact on neurodevelopment and confer risk for psychiatric disorders, we investigated the effect of a loss of *NT5C2* function in these cells at the transcriptomic and molecular level. Specifically, we knocked down expression of *NT5C2* in hNPCs using an RNAi approach. CTX0E16 hNPCs were transfected using two independent, non-overlapping small interfering RNA (siRNA) sequences, A and B. The efficacy of the siRNA transfection was determined by uptake of a fluorescently labelled oligonucleotide (BLOCK-iT), which revealed a transfection rate of 90% ± 0.02, relative to the total number of cells (n = 4 biological replicates per condition; **Figure 3A**). Next, we assessed the extent of the knockdown on *NT5C2* expression elicited by a 72-hour incubation with siRNAs A and B, relative to the control cultures treated with a scramble sequence (**Figure 3B**). This analysis revealed that *NT5C2* expression was significantly affected by siRNA treatment (One-way ANOVA, F (2, 8) = 13.45, P = .003). Tukey post hoc tests revealed a mean ∼27% decrease in *NT5C2* expression in knockdown conditions (siRNA A: 71.00 ± 13.92, P = .004; siRNA B: 75.10 ± 7.29, P = .004). We assessed the ability of the siRNAs to knockdown the NT5C2 protein in independent hNPC cultures (**Figure 3C-D**). This analysis revealed a significant decrease in abundance of this protein in the knockdown conditions (One-way ANOVA, F (2, 41) = 12.23, P < .001; Tukey post hoc tests: siRNA A, 58.79 ± 34.74, P < .001; siRNA B, 62.42 ± 21.54, P < .001).

**Figure 3.**
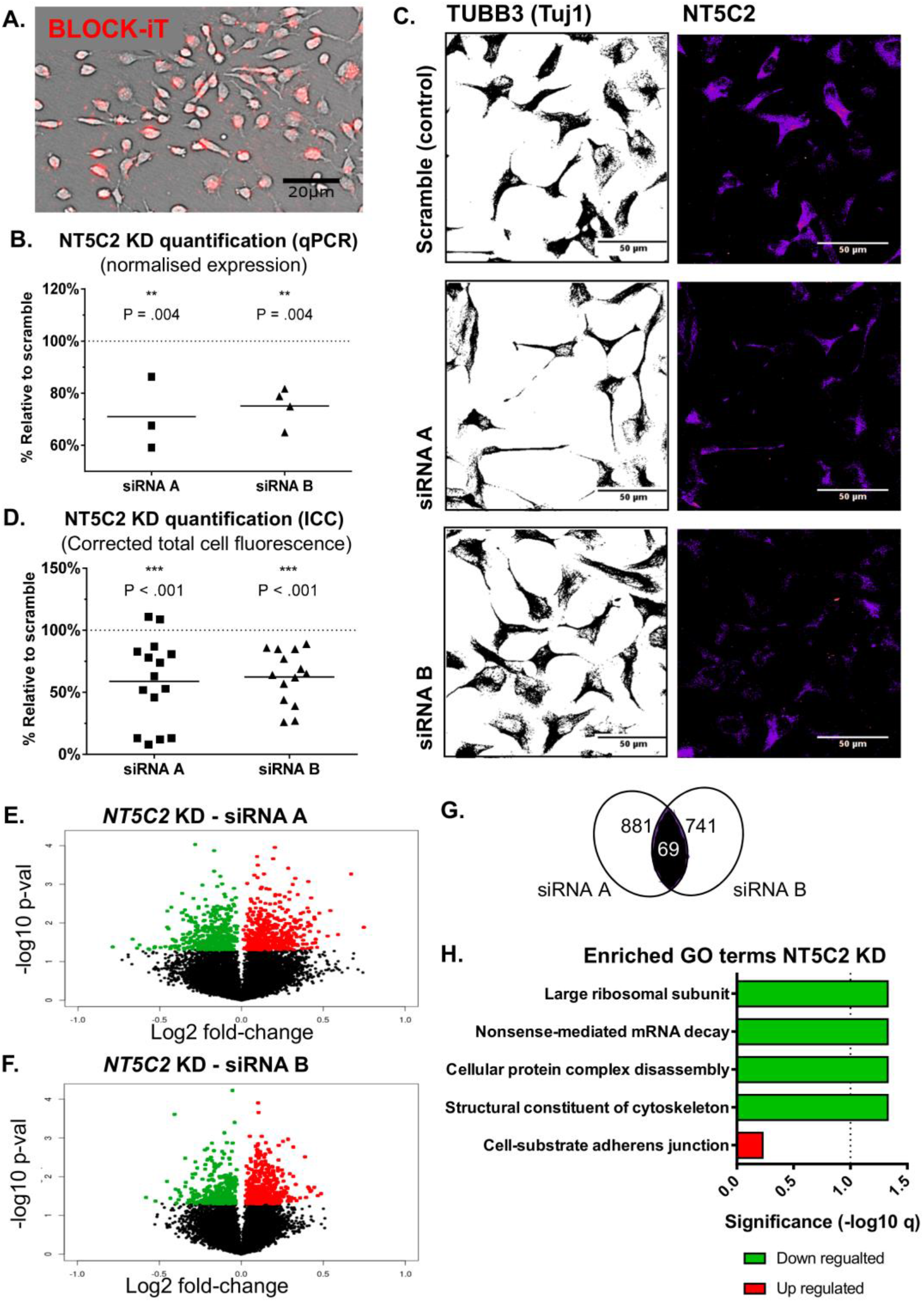
Knockdown of *NT5C2* in hNPCs elicits transcriptomic changes associated with protein translation. **(A)** The efficacy of the siRNA transfection was determined by uptake of BLOCK-iT, a fluorescently labelled oligonucleotide. Over 90% of cells contained this reagent after incubation with this reagent for 72 hrs. **(B)** NT5C2 expression was significantly reduced in knockdown cultures (one-way ANOVA, Tukey post hoc tests, **P <.01). **(C, D)** The ability of the siRNA treatments to significantly reduce NT5C2 expression in hNPCs, at the protein level (one-way ANOVA, Tukey post hoc tests, ***P < .001). **(E)** Volcano plots indicating nominally significant transcriptomic changes elicited by siRNA A, and **(F)** siRNA B; **(G)** Venn diagrams indicates number of common genes differentially regulated by siRNA A and B. The overlap is significant according to a Fisher’s exact test (P < .001). **(H)** Gene ontology terms enriched within the genes concordantly and differentially expressed in both knockdown conditions. The line indicates the significance threshold (−log10 (q < .05)).

Transcriptomic analysis of the knockdown samples revealed that *NT5C2* siRNAs A and B elicited expression changes to 881 and 741 genes, respectively (linear regression, nominal P < .05; **Figures 3E** and **F**). To reduce off-target effects associated with individual siRNAs^34^, we identified the concordant transcriptomic effects *shared* between both siRNA treatments. This analysis revealed an overlap of 69 genes (**Figure 3G**), which is unlikely to occur by chance given the number of genes tagged in the microarray (n = 21,196 genes; Fisher’s exact test, P < .001, Jaccard index < .001, odds ratio = 2.6; gene list in **Supplementary Table 13**). This list was subdivided by directionality of effect, and the up- and downregulated network topologies were calculated using GeneMania^28^ (**Supplementary Figure 5**). This analysis revealed multiple edges that indicate co-expression and co-localisation of genes/nodes in the networks, corroborating their functional association. The up- and downregulated gene lists were further analysed for enrichment of gene ontology (GO) terms, which revealed significantly downregulated terms (FDR < .05) pertaining to the regulation of protein translation (GO:0015934 large ribosomal subunit; GO:0043624 cellular protein complex disassembly; GO:0000184 nonsense-mediated mRNA decay) and of the cytoskeleton (GO:0005200 structural constituent of the cytoskeleton; **Figure 3H**, **Supplementary Table 14**). The top upregulated GO term suggested the involvement of *NT5C2* in cell adhesion (GO:0005924 cell-substrate adherens junction), but this term did not survive multiple testing correction (q > .05). We used RT-qPCR to validate a panel of gene expression changes detected in the microarray analysis which are functionally related to the GO terms, to support their association with *NT5C2* (**Supplementary Figure 6**). These included changes to the heterogeneous nuclear ribonucleoprotein A1 (*HNRNPA1)*, which is implicated in protein translation^35^; the proteasome 26S subunit, ATPase 4 (*PSMC4)*, involved in protein degradation, cytoskeleton remodelling^36^, and Parkinson’s disease^37^; and the autophagy-related cysteine peptidase gene (*ATG4B)*, involved in cytoskeleton regulation^38^. Altogether, these findings support a risk mechanism for psychiatric disorders in hNPCs, in which decreased *NT5C2* expression causes global changes to protein translation.

### *NT5C2* regulates AMPK and ribosomal protein S6 (rpS6) phosphorylation

Protein translation has been extensively implicated in psychiatric disorders^39-41^, but the mechanisms leading to disruptions in this process remain elusive. As our transcriptomic data indicated that reducing *NT5C2* expression could impact translation machinery, we were interested in investing whether this could be associated with an alteration in protein translation. One mechanism via which *NT5C2* expression may impact protein translation is through AMPK signalling. For example, in glioblastoma cells, NT5C2 has been shown to regulate this signalling cascade^19^. Interestingly, in neurons, AMPK is part of a signalling cascade that links extra cellular signals, including synaptic-activity, with the regulation of ribosomal activity, and thus protein synthesis under physiological conditions^42^. Altered AMPK signalling in neurodevelopmental disorders as well as in Alzheimer’s Disease pathophysiology has also been linked with abnormal regulation of protein synthesis^43, 44^. Therefore, we tested whether knockdown of NT5C2 resulted in abnormal AMPK signalling and in altered protein translation in hNPCs. We observed a significant effect of the *NT5C2* knockdown on total AMPK-alpha expression (Kruskal-Wallis test, H(3) = 12.23, P < .001); a mean 132% increase in the abundance of this kinase (Dunn’s post hoc tests: siRNA A, Mdn = 236.10, P = .002; siRNA B, Mdn = 182.8, P = .017; **Figure 4A**). In addition, we observed a significant effect of the knockdown on the level of phosphorylated AMPK-alpha (Thr172) (Kruskal-Wallis test, H(3) = 7.65, P < .013). The knockdown elicited a mean 55% increase in phosphorylated AMPK relative to control cultures (Dunn’s post hoc tests: siRNA A, Mdn = 141.2, P = .033; siRNA B, Mdn = 160.7, P = .033; **Figure 4A**). These findings corroborate a functional association between *NT5C2* and AMPK signalling in hNPCs, and further suggest that this kinase pathway may be dysregulated in psychiatric disorders.

**Figure 4.**
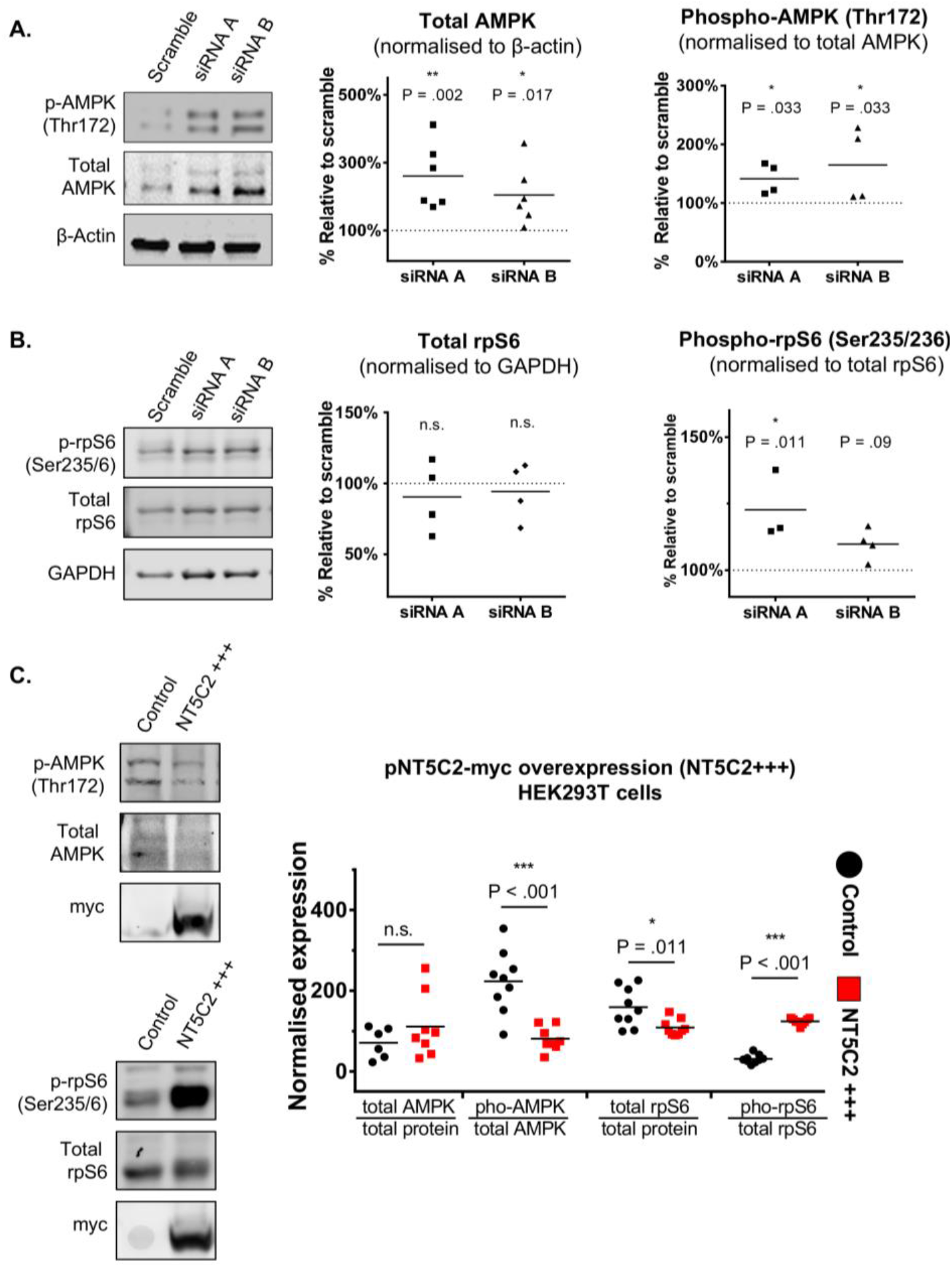
NT5C2 regulates AMPK signalling and rpS6 activity. **(A)** *NT5C2* knockdown elicited a mean 132% increase in total AMPK-alpha levels (Kruskal-Wallis test, Dunn’s post hoc tests, **P < .01, *P < .05), and in phosphorylated AMPK-alpha (Thr172) in hNPCs (*P <.05). **(B)** Knockdown of *NT5C2* did not elicit changes to total rpS6 in hNPCs (P > .05); there was, however, an effect on phosphorylated rpS6 (Ser235/Ser236) levels, whereby siRNA A elicited a mean 23% increase (Kruskal-Wallis test, Dunn’s test, *P < .05), and siRNA B a modest mean 10% increase, which was not significant after correction (Dunn’s test, P = .09). **(C)** The overexpression of *NT5C2* in HEK293T cells causes a significant decrease in phosphorylated AMPK-alpha levels and in total rpS6, and a significant increase in phosphorylated rpS6 (independent t-test, ***P < .001, *P = .01).

As AMPK signalling has been linked with the regulation of protein synthesis^42-44^, and owing to our transcriptomic data, we next tested whether the *NT5C2* knockdown had an effect on the phosphorylation of the ribosomal protein S6 (rpS6) at residues Ser235/Ser236. Assessment of rpS6 phosphorylation is widely used to monitor activation of *mammalian target of rapamycin complex 1* (mTORC1) signalling and can be used as a proxy to estimate protein translation in neurons^45^. Knockdown of *NT5C2* did not alter levels of total rpS6 levels (Kruskal-Wallis test, H(3) = 0.04, P > .05; **Figure 4B**). However, a significant increase in rpS6 phosphorylation was observed in knockdown conditions compared to the scramble-treated controls (Kruskal-Wallis test, H(3) = 8.22, P = .002). *NT5C2* knockdown elicited by siRNA A was significantly associated with a mean 23% increase in phosphorylated rpS6 (Dunn’s post hoc test, Mdn = 115.9, P = .012; **Figure 4B**), whilst the knockdown with siRNA B elicited a mean 10% increase (Dunn’s test, Mdn = 110.20, P = .09; **Figure 4B**). These data suggest that *NT5C2* is a negative regulator of rpS6 phosphorylation and thus protein translation, in hNPCs.

To support the association between *NT5C2*, AMPK signalling and rpS6 activity, we carried out complementary experiments in HEK293T cells using a pNT5C2-myc overexpression plasmid. Exogenous expression of *NT5C2* in HEK293T cells resulted in a mean ∼64% decrease in phosphorylated AMPK-alpha (independent t-test; control: 223.00 ± 76.99, overexpression: 81.05 ± 30.14, t(15) = 4.88, P < .001, Bonferroni corrected P < .001; **Figure 4C**), whilst no difference in total AMPK levels were observed (t(12) = 1.16, P > .05; **Figure 4C**). These data are consistent with our data indicating that NT5C2 is a negative regulator of AMPK signalling. Next, we examined levels of total and phosphorylated rpS6 in the presence or absence of ectopic NT5C2. This revealed a mean ∼28% decrease in total rpS6 abundance (control: 159.10 ± 48.52, overexpression: 108.8 ± 48.52, t(16) = 2.88, P = .011, corrected P = .044; **Figure 4C**), and a mean 300% increase in rpS6 phosphorylation (control: 31.03 ± 10.66, overexpression: 124.10 ± 8.20, t(16) = 20.76, P < .001, corrected P < .001; **Figure 4C**). This effect of exogenous *NT5C2* on rpS6 phosphorylation was opposite to that observed in hNPCs, corroborating the complex nature of the intracellular cascades governing protein translation. Moreover, this could reflect different regulatory systems regulating protein translation in the different cell types. Ultimately, these data demonstrate that NT5C2 regulates the function of components involved in the regulation of protein translation. These data suggest that risk alleles which decrease expression of *NT5C2* may confer risk to psychiatric disorders via changes to protein translation.

### Knockdown of the *NT5C2* homologue *CG32549* in *D. melanogaster* is associated with abnormal climbing behaviour

Our *NT5C2* knockdown studies in hNPCs, together with the developmental profile of the gene in human brain (**Figure 2A**) support an important role for *NT5C2* in early brain development. However, these molecular studies do not afford an insight into the potential impact of reduced developmental expression of *NT5C2* at a systems level. Interestingly, the NT5C2 protein shares 60.5% sequence identity and 80.2% sequence similarity with the *D. melanogaster* homologue, *CG32549* (**Supplementary Figure 7**), suggesting that they likely exert the same or similar function. Thus, we reasoned that it would be possible to gain an insight into the functional impact of reduced *NT5C2* expression *in vivo*, by modelling the knockdown of *CG32549* on a complex and polygenic behaviour, such as climbing. This is a polygenic psychomotor trait driven by an interaction between cognitive function and physical activity, which is easily observable and measurable in *D. melanogaster*, and that has been previously used to study the functional impact of gene mutations in *D. melanogaster* ^29, 46^, including those associated with psychiatric and neurodegenerative risk^47^.

To test the role of *CG32549 in vivo*, three knockdown fly lines were engineered using the Gal4 upstream activating sequence (UAS-Gal4) system^48^, to artificially reduce *CG32549* expression either ubiquitously throughout the whole body, specifically in the central nervous system (CNS), or in the gut. This was achieved by crossing a *CG32549*-RNAi line with flies containing the GAL4-driven promoters of *ACT5C*, *ELAV*, or *GUT*, respectively (**Figure 5A**). The Gal4-UAS system is most active at 29°C^49^, and therefore crosses were incubated at this temperature from the pupal stage to elicit the strongest knockdown of *CG32549* at this point of development, when new neurons are still being formed^50^. Importantly, this allowed us to disentangle the potential function of *CG32549* in non-neuronal cells, such as muscles, which may influence performance in the negative geotaxis assay. The ubiquitous knockdown of *CG32549* was associated with reduced expression of this gene by 87% in the brain of flies with the ubiquitous knockdown (median, Mdn = 0.19) relative to flies without the RNAi cassette (Mdn = 1.00; Mann-Whitney test, two-tailed, U < .001, P = .029, n = 4; Figure 2A). No difference in survival was observed across the different genotypes (Mann-Whitney test, P > .05, n = 6; **Figure 5C**). There was, however, a mean ∼20% reduction in climbing success in flies with ubiquitous (Mdn = 0.78) and neuronal-specific knockdowns (Mdn = 0.88), relative to flies without the RNAi cassette (Mdn = 1.00; Mann-Whitney tests, U < .001, P = .002 for both comparisons, Bonferroni corrected P-values = .006, n = 6; **Figure 5C**). This effect was not observed in flies with knockdown of *CG32549* restricted to gut (Mann-Whitney test, P > .05, n = 6; **Figure 5C**). There was no difference in climbing impairment observed between flies with ubiquitous (80.00% ± 7.31) or neuronal-specific knockdowns (85.41% ± 5.30; One-way ANOVA, F (2, 15) = 16.58, P < .001, n = 6 per condition; Tukey post hoc test P > .05), although these effects significantly differed from that observed in flies with the gut-specific knockdown (97.62% ± 2.64; gut vs. ubiquitous: P <.001; gut vs. neuronal: P = .004). Taken together, these results demonstrate the function of NT5C2 at a systems level, implicating it in a complex, measurable phenotype in *D. melanogaster*.

**Figure 5.**
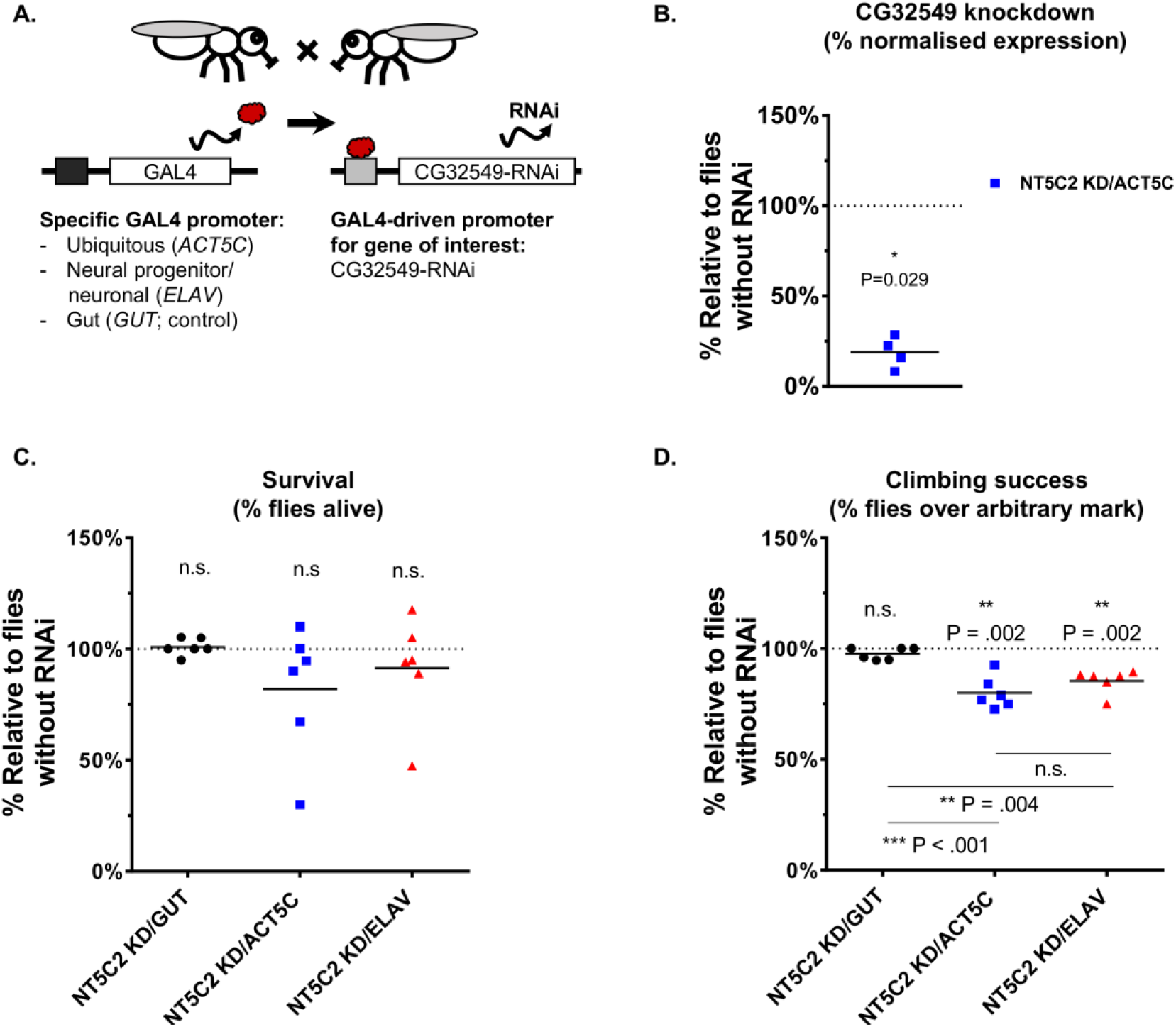
The knockdown of *CG32549 (NT5C2 homologue)* in *D. melanogaster* elicited by RNAi directed ubiquitously (ACT5C), in gut (GUT), or in neural progenitor cells/neurons (ELAV). **(A)** Schematic representation of the knockdown using the GAL4/UAS system. **(B)** *CG32549* was less expressed in the brain of knockdown flies (Mann-Whitney test, *P < .05). **(C)** The knockdown did not significantly affect survival measured 17-20 days after eclosure (Mann-Whitney test, P > .05). **(D)** Climbing impairment observed in flies where knockdown occurred ubiquitously or in neurons (Mann-Whitney tests, **P < .01), but not in gut (P > .05). There was no difference in the climbing impairments observed between flies with ubiquitous or neuronal-specific knockdowns (one-way ANOVA, Tukey post hoc test, P > .05), although these effects significantly differed from that observed in flies with the gut-specific knockdown (Tukey tests: gut vs. ubiquitous: ***P < .001; gut vs. neuronal: **P < .01)

## Discussion

This study aimed to investigate the expression and function of *NT5C2* in the adult brain and in neural progenitor cells, and to characterise the biological mechanisms via which this gene governs risk for psychiatric disorders. Previously, we identified cis-regulatory mechanisms elicited by common psychiatric risk alleles on chromosome 10q24 reducing expression of *NT5C2* in the hippocampus, dorsolateral prefrontal cortex and nucleus caudate of population controls, and in the second trimester foetal brain^3^. Here, we observe that *NT5C2* expression is indeed decreased in the hippocampus of psychiatric patients, and our data from validated human cellular assays indicate that this may result in changes to protein translation in neurons. Whilst the effect of *NT5C2* on protein translation was observed in hNPCs (Figure 5), where this protein is highly expressed and ubiquitously distributed (Figure 3), we hypothesise that this mechanism may also occur in the adult brain, although it may be susceptible to additional regulatory mechanisms. We further hypothesise that the psychiatric risk mechanism pertaining *NT5C2* expression in the adult brain is more likely to occur in neurons, where this protein is more expressed relative to glial cells (Figure 1). This is consistent with the recent suggestion that expression of psychiatric risk genes is cell-type specific, with particular enrichment of neuronal cell type^32^. Ultimately, the association between *NT5C2* function and protein translation regulation corroborates the idea that this gene serves a fundamental role in cell biology, as it is implicated in a multitude of disease states^1, 2, 4-9^.

The transcriptomic analysis of the knockdown in hNPCs also revealed that *NT5C2* may govern genes that regulate the cytoskeleton (Figure 4). Changes to the cytoskeleton have been previously implicated in psychiatric disorders^51^, but these significantly depend on the translational machinery^52^. Protein translation is such a finely regulated process; it is in part modulated by AMPK signalling^53-55^ and rpS6 activity^45, 56, 57^, and has been vastly implicated in psychiatric disorders^39-41^. Interestingly, AMPK has been linked to neuroprotection and aspects of neurodevelopment that are relevant to psychiatric disorders, such as axogenesis and bioenergetics^58-60^. rpS6 activity, in turn, has been associated with increased neuronal function, neuroplasticity, and modulation by antipsychotics (e.g. clozapine, haloperidol, olanzapine) and abuse drugs (e.g. cocaine, methamphetamine, and tetrahydrocannabinol)^45^. While it is possible that *NT5C2* modulates protein translation by binding directly to other members of the translation machinery^61^, our data suggest that AMPK signalling and rpS6 activity could be explored as tractable drug targets for psychiatric disorders. The knockdown of the fly homologue *CG32549* in Drosophila further implicates neuronal *NT5C2* expression in motility behaviour (**Figure 5**), corroborating the importance of this gene at a systems level. It is possible that this effect in motility is driven by changes to the rate of protein translation, as it has been demonstrated that abnormal AMPK and rpS6 activation in Drosophila are associated with climbing impairments^62, 63^.

There are two main limitations to our study which should be acknowledged. First, our gene expression analysis of case-control differences is underpowered, especially when considering the heterogeneity inherent to all psychiatric conditions. We partly addressed this issue by controlling for gene expression changes associated with the effect of demographics; this analysis could be further improved by significantly increasing sample size. Second, we obtained a modest knockdown in the loss-of-function experiment in hNPCs, which is likely due to the proliferative nature of these cells. We partly addressed this issue by testing hypotheses generated based on the microarray results using western blotting and a different cell type (HEK293T cells; **Figure 4**), which supported our findings.

Our results provide clues to the mechanism via which genetic variation affecting *NT5C2* expression may confer risk for psychiatric disorders. While functional studies *in vitro* or using model organisms cannot entirely capture the complex nature of these conditions, we anticipate that further work on this and other susceptibility genes may lead to the identification of converging risk mechanisms, which may reveal novel drug targets amenable for therapeutic intervention, or biomarkers for these conditions.

## Acknowledgements

This work was supported by CAPES (Ministry of Education, Brazil, award no. BEX1279-13-0 to RRRD), an NIHR Maudsley Biomedical Research Centre Career Development Award (RRRD); the Medical Research Council UK (G0802166 to NJB; Skills Development Fellowship MR/N014863/1 to TRP; MR/N025377/1 to ACV; MR/L021064/1 to DPS) and MRC Centre grant (MR/N026063/1); the Wellcome Trust ISSF (097819 to DPS); the King’s Health Partners Research and Development Challenge Fund, a fund administered on behalf of King’s Health Partners by Guy’s and St Thomas’ Charity, Royal Society UK, Brain & Behavior Foundation (formally National Alliance for Research on Schizophrenia and Depression, NARSAD), the Psychiatric Research Trust (DPS); the European Union’s Seventh Framework Programme (FP7-HEALTH-603016) (DPS); and US National Institutes of Health grants CA206488 and AI076059 (DFN). This study represents independent research part funded by the NIHR-Wellcome Trust King’s Clinical Research Facility and the National Institute for Health Research (NIHR) Biomedical Research Centre at South London and Maudsley NHS Foundation Trust and King’s College London. The views expressed are those of the authors and not necessarily those of the NHS, the NIHR or the Department of Health and Social Care. Tissue samples were supplied by The London Neurodegenerative Diseases Brain Bank, which receives funding from the MRC and as part of the Brains for Dementia Research programme, jointly funded by Alzheimer’s Research UK and Alzheimer’s Society. We thank Dr. Frank Hirth for constructive reading of the manuscript.

## Authorship contribution

Conceived and designed experiments: RRRD, DPS, NJB. Performed the experiments: RRRD, NDB, GAH, MCC. Analysed the data: RRRD, TRP. Contributed reagents, biological material, expertise: SHL, SS, IAW, CT, GDB, ACV, IE, DFN, RMM, NJB, TRP. Wrote the paper: RRRD, DPS.

